# Evidence of two centromeres in the germline-restricted chromosome (GRC): insights from zebra finch lampbrush chromosomes

**DOI:** 10.64898/2026.07.12.738116

**Authors:** Olga Takki, Valeriia Volodkina, Nikolay Rubtsov, Kira Zadesenets, Francisco J. Ruiz-Ruano, Niki Vontzou, Julia Jukova, Maria Kulak, Elena Gaginskaya, Alexander Suh, Svetlana Galkina

**Author notes:** Institute of Ecology and Evolution, The University of Edinburgh, Edinburgh, EH9 3FL, UK.

## Abstract

The germline-restricted chromosome (GRC) of the zebra finch *Taeniopygia guttata* represents a well-established model of programmed DNA elimination in vertebrates. Although the DNA composition of the GRC, as well as elimination processes during spermatogenesis and early embryogenesis, have been characterised previously, little is known about the cytogenetic features underlying its unusual behaviour, including its stable transmission through the maternal germline. Here, we provide a detailed characterisation of the zebra finch GRC at the diplotene stage of female meiosis, when chromosomes are actively transcribed and acquire the form of giant lampbrushes. We identified a transcriptionally repressed region on the GRC, which we term the ’belt’. Microdissection and sequencing of the belt revealed that it is predominantly composed of a tandem repeat derived from the *dph6* gene, *robo1* gene fragments, and ERVs. Notably, the terminally located functional centromere of the GRC lacks typical zebra finch centromeric satellites and, conversely, consists of the newly identified GRC-specific tandem repeats *Tgut16-201* and *Tgut17-167.* The canonical centromeric repeat *Tgut716* was observed in the GRC belts. Moreover, belts, like the terminal GRC centromere, were associated with coilin-containing nuclear bodies, which serve as markers of centromeric regions on zebra finch lampbrush chromosomes. Together, our findings provide evidence for the presence of one functional and one putative centromeric region on the zebra finch GRC, suggesting their role in non-Mendelian inheritance of the GRC.

**Author summary:** Germline-restricted chromosomes (GRCs) are unusual chromosomes that are retained in germ cells but eliminated from somatic cells during early development. They have evolved independently in several groups of organisms, but are particularly notable in passerine birds, a large monophyletic vertebrate clade (∼6,700 species) in which GRCs have persisted for at least 44 million years.

Passerine GRCs are normally transmitted to the next generation through the maternal germ cell, however, the mechanisms ensuring their inheritance remain unknown. To address this question, we examined the structure of the zebra finch GRC during female meiosis. We found that the GRC differs from all other chromosomes in possessing two distinct centromeric regions: a functional terminal centromere and an extended heterochromatic region exhibiting centromeric properties. These unusual features suggest a mechanism by which the GRC may achieve its preferential transmission through the female germline.

Our findings substantially advance the understanding of the zebra finch GRC and the general biology of passerine GRCs. By revealing chromosome features that may underlie their non-Mendelian inheritance, this work provides new insights into the evolution and behaviour of GRCs and other selfish chromosomes that bias their own transmission.

## Introduction

The discovery of the germline-restricted chromosome (GRC) in the germline karyotypes of songbirds [1–3] represents a major advance in avian cytogenetics as an example of programmed genome elimination among vertebrates [4]. The widespread detection of the GRC across diverse passerine lineages [2,5–12] supports the hypothesis of an ancient GRC origin from an autosome in the common ancestor of passerine approximately 44 million years ago, predating the suboscine-oscine divergence [12]. Passerine GRCs show striking interspecific variation in size and in their genetic content, reflecting dynamic evolutionary history [2,3,13]. One of the most intriguing aspects of the GRC is its non-Mendelian mode of inheritance. In germ cells, it typically exists in a single copy in males and two copies in females [1–3]. However, in some cases the copy number of the GRC can deviate from the typical univalent (male) and bivalent (female) states. These deviations, which include variation in GRC copy number within and between individuals, rare cases of paternal transmission, and elimination of the GRC from somatic cells, are likely the phenotypic result of segregation instability [1,3,14–17].

The zebra finch (*Taeniopygia guttata*, Estrildidae, Passeriformes) is the species in which the GRC was first discovered in birds [1], and is at the forefront of GRC research. This status has been strengthened by the availability of a high-quality genome assembly of this well-established model organism [18–20], which enables comprehensive comparisons between the genomes of somatic and germ cells [21,22]. In the zebra finch, the GRC is the largest chromosome in germ cells [1], comprising over 10% of the genome and spanning approximately 160 Mbp [21]. At least 125 different protein-coding genes have been identified in the zebra finch GRC [12], the majority of which are paralogs of genes located in the autosomes and the Z-chromosome [21,22]. These genes are expressed throughout gametogenesis, from early embryogenesis in the unlaid egg to adult ovaries and testes [21–24]. Functional annotation suggests that several GRC-linked genes may contribute to germline specification (e.g., *itga11_GRC_, tfeb_GRC_, scrib_GRC_*) and female gonadal development (e.g. *bmp15_GRC_, zglp1_GRC_, itga11_GRC_*), whereas the functions of others, including genes involved in cell cycle regulation genes (e.g. *pbk_GRC_, cpeb1_GRC_*), remain less understood [23]. In adult birds, GRC-linked gene expression is strongly biased toward the ovary, with substantially lower transcription activity in the testis.

Consistent with this expression profile, the univalent state of the zebra finch GRC in spermatocytes is thought to underlie its elimination during the first meiotic division. During pachytene, the GRC undergoes progressive heterochromatinisation (marked by the accumulation of H3K9me2 and H3K9me3), acquires double-strand breaks, and condenses into a compact DAPI-positive body that is expelled from the nucleus [5,25,26]. Its elimination is further facilitated by the absence of the inner centromere protein INCENP, which causes delayed spindle attachment and subsequent arrest at the metaphase equator [25]. The expelled GRC subsequently forms a micronucleus that is ejected from the cell [1,5,14,25].

In contrast to spermatocytes, the zebra finch GRC is present as a bivalent in oocytes. At the pachytene stage of female meiosis, the GRC has been described as an acrocentric chromosome [1,14,25] and its bivalent typically carrying two or occasionally three MLH1-positive recombination nodules [2,14]. However, its organisation at subsequent stages of female meiosis remains unknown, including the diplotene stage of prophase I. During diplotene, avian chromosomes transform into lampbrush chromosomes (LBCs), in which extensive transcriptional activity gives rise to lateral loops enriched in ribonucleoprotein (RNP) complexes [27]. The approximately 30-fold increase in size relative to metaphase [28,29] makes LBCs an exceptionally powerful system for investigating chromosome architecture. High level of transcriptional activity, together with the distinctive chromomere-loop pattern, enables high-resolution cytogenetic mapping using fluorescence *in situ* hybridisation (FISH), facilitating precise localisation of sequences, construction of detailed cytogenetic maps, and identification of transcriptionally active and silent regions. LBCs have proven particularly valuable for analysing regions that are difficult to investigate in their condensed state, such as the W chromosome [30–35] and centromeric regions [36–39].

Previously, we identified the zebra finch GRC bivalent at the lampbrush stage using FISH with a microdissected probe [2]. RNA polymerase II immunolabelling revealed transcriptional activity across the GRC, comparable to that of other LBCs, except for a prominent DAPI-positive region. These observations provided an entry point for investigating the structural features of the GRC that may underlie its unusual inheritance, the primary focus of the present study. We performed high-resolution cytological mapping of the zebra finch GRC using a panel of molecular markers, including newly identified GRC-specific tandem repeats. We also investigated the molecular composition of prominent DAPI-positive lampbrush GRC regions, termed the “belts”. By integrating our data with previously reported GRC markers, we constructed a comprehensive cytological map of the chromosome. We propose that the peculiar structural organisation of the GRC can explain its non-Mendelian inheritance through the maternal germline.

## Results

### Zebra Finch Lampbrush Chromosomes: GRC

The general morphology of zebra finch LBCs differs from that previously described for other birds. We did not observe distinct morphological features, such as prominent interstitial marker loops, that would allow for the consistent identification of individual chromosomes (Fig 1). Instead, the chromosomal set is largely homogeneous, with most chromosomes exhibiting simple lateral loops. Each pair of homologous chromosomes forms bivalent, held together by one or two chiasmata. The latter are typically located near both ends of the homologues, so the bivalent spread on the slide can take the form of a ring (Fig 1). A common feature of all chromosomes is the association with the characteristic RNP complexes at the ends of bivalents, terminal giant loops (TGLs), and intranuclear bodies. These latter are referred to as nuclear biomolecular condensates (NBioMCs), which typically associate with the centromeric regions of LBCs [40]. For the macrochromosomes, their identification was only possible by relative size, while we recognised the sex ZW bivalent by its asymmetry and the bright DAPI-positive fluorescence of the heterochromatic W chromosome (Fig 1a’).

**Figure 1.**
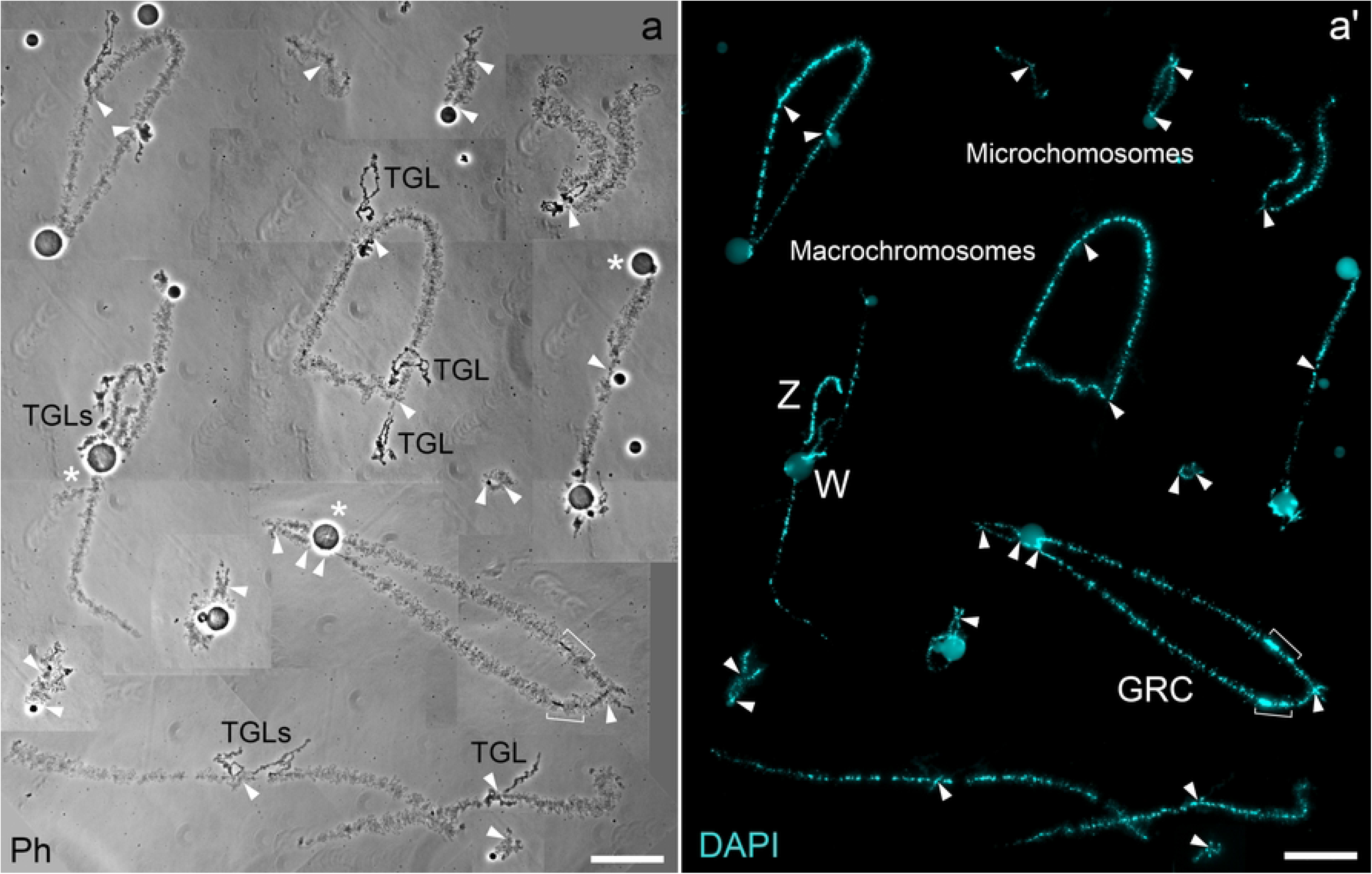
Part of a chromosome set obtained microsurgically from a zebra finch previtellogenic oocyte (the full set consists of 41 bivalents). (a) Several LBCs visualised using the phase contrast. (a’) The same chromosomes are stained with DAPI. Macrochromosomes and microchromosomes are visible, homologous chromosomes are united by chiasmata (arrowheads) into bivalents. Homologues may possess terminal giant loops (TGLs in the a), which are DAPI-negative (invisible in the a’). Nuclear biomolecular condensates (NBioMCs) indicated by asterisk are observed either associated with bivalents or free within the nucleoplasm. The Z and W chromosomes are distinguished by their pattern of heterochromatinization causing asymmetrical bivalent morphology; the GRC by its loopless regions, the “belts” (square brackets). Scale bars: 20 μm.

Against this uniform background, the GRC bivalent is strikingly different and easily identifiable. It is the largest bivalent in the karyotype, reaching a maximum length of approximately 250 μm. Each of its semi-bivalents can consist of around 130 DAPI-positive chromomeres at peak decondensation. We consistently observed only two terminal chiasmata located at both ends of the GRC bivalent (Fig 1 and 2). A defining characteristic of the GRC is the presence of condensed, DAPI-positive, loopless regions on both homologues, which we term “belts”. These belts are transcriptionally silent, as evidenced by the absence of lateral loops (Fig 1 and 2), enrichment of 5-methylcytosine (Fig 2a’’), and the previously documented absence of RNA polymerase II [2]. The GRC at the lampbrush stage exhibits two distinct regions potentially associated with NBioMCs. The first is the GRC end attached to the large (about 6 µm in diameter), spherical, light-refracting structure visible under phase contrast (Fig 2b) and confirmed to be coilin-positive (Fig 2b’’). The second are loopless belts, associated with numerous smaller bodies (< 0.5 µm in diameter) of a similar nature (Fig 2b), although their presence is not an obligatory feature of the belts.

**Figure 2.**
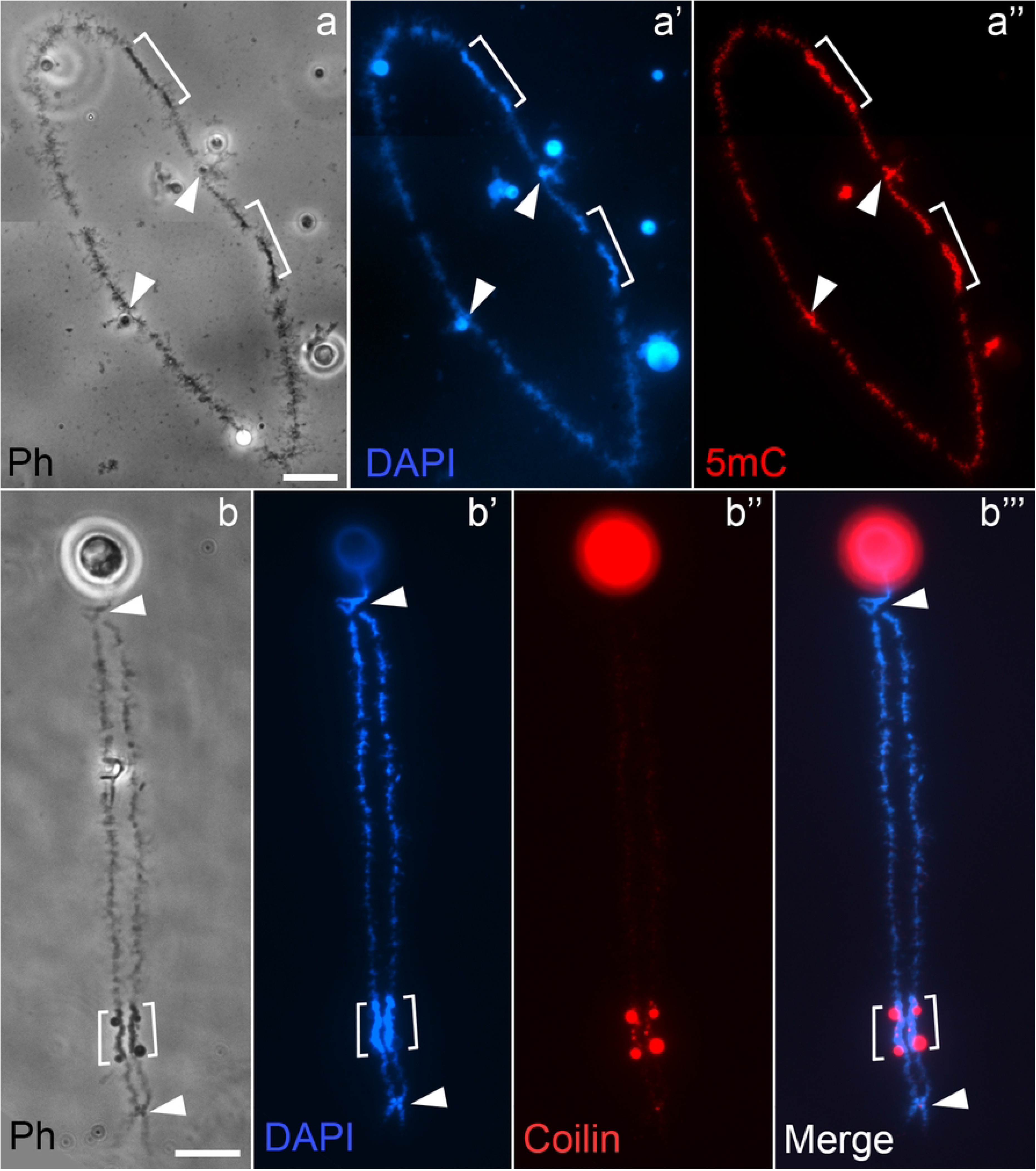
Immunostaining of the lampbrush GRC with antibodies against 5-methylcytosine (a-a’’) and coilin (b-b’’’). Enrichment of 5-methylcytosine (5mC) (red fluorescence) is observed along the GRC axis, including the belt region (a’’). Antibodies against coilin (red fluorescence) mark intranuclear bodies or NBioMCs: a large body is located at one chromosomal end, while smaller bodies are often associated with the belt region near the opposite end. Arrowheads indicate chiasmata, square brackets denote the belt region. Scale bars: 10 µm.

### GRC Belt Sequence Composition

The transcriptional silencing of the GRC belts prompted us to precisely study their composition belts. Therefore, we performed chromosome microdissection of the GRC belt region to obtain material for Illumina library preparation and sequencing. To test the specificity of the GRC belt composition, a FISH probe was generated from the microdissected material (Fig 3a). Hybridisation with the obtained probe produced a signal exclusively in the belt regions on both the lampbrush- and pachytene-stage GRCs (S1a and S1c Figs). Furthermore, the probe specifically labelled micronuclei containing the eliminating chromatin in testicular cell preparations, confirming its GRC specificity (S1b Fig).

**Figure 3.**
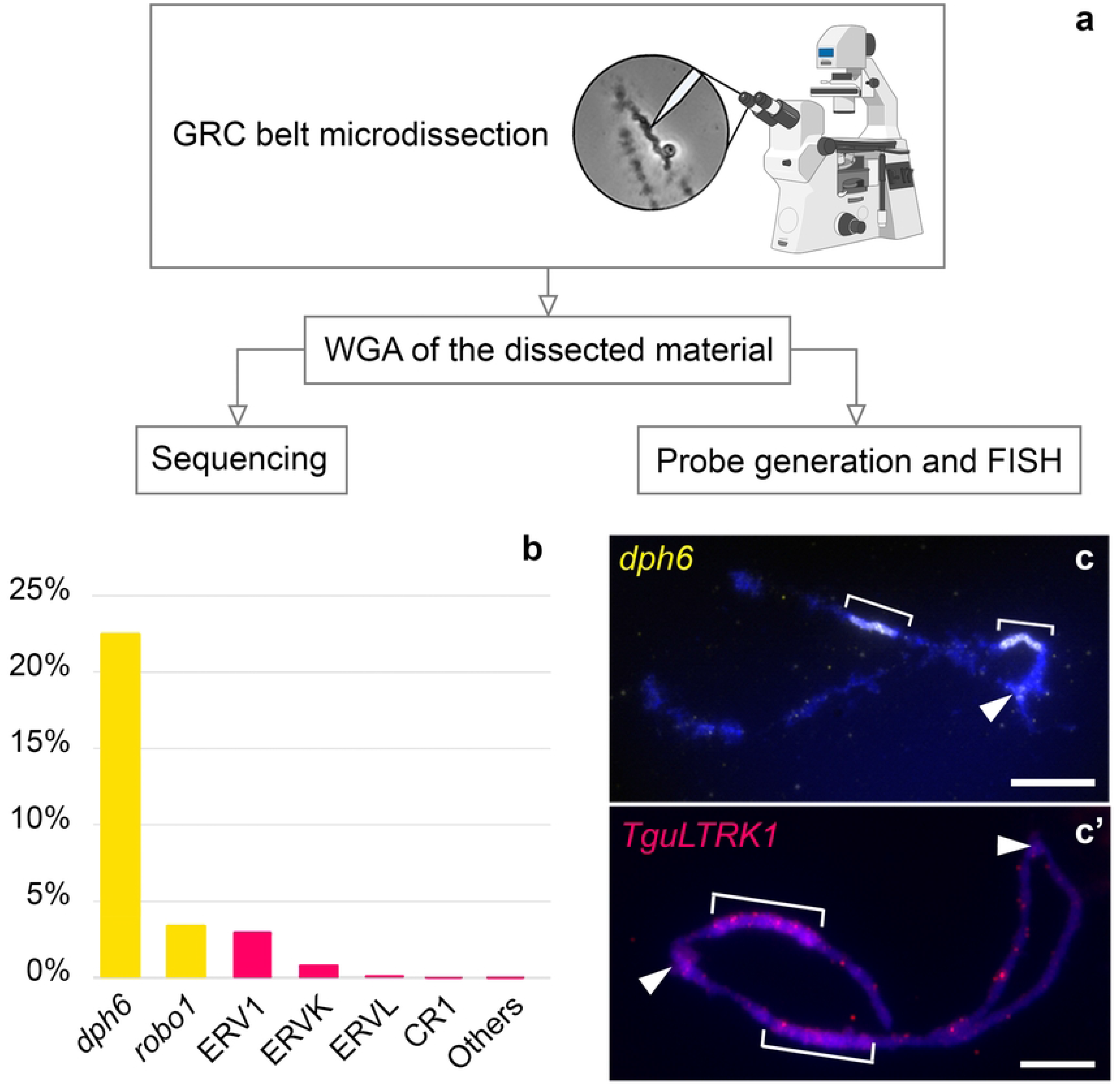
Analysis of the zebra finch GRC belt region. (a) Schematic overview of the experimental workflow: microdissection of the GRC belt region and whole-genome amplification (WGA), followed by preparation of a FISH probe and a sequencing library, and downstream FISH analysis and Illumina sequencing. (b) Repeat composition of the microdissected GRC belt as revealed by the Illumina library. The y-axis represents the proportion of each repeat in the library. Yellow bars indicate repeats derived from single-copy genes (*dph6*, *robo1*) and red bars indicate transposable elements (ERV1, ERVK, etc.). (c, c’) Hybridisation results with the dph6 probe (yellow fluorescence, c) and the TguLTRK1 probe (red fluorescence, c’). Chromosomes are counterstained with DAPI (blue fluorescence). Arrowheads indicate chiasmata, square brackets denote the belt region. Scale bars: 10 µm.

Sequencing of the microdissected material resulted in the library, 26.14% of which was mapped to the zebra finch somatic reference genome (bTaeGut1.4.pri): 18.30% aligned to two fragments of the *dph6* gene (20.4 kb and 5.75 kb segments), while 1.87% mapped to a 2.5 Mb region on chromosome 1 containing the *robo1* and *gbe1* genes (S1 Table). For more detailed repetitive elements characterisation, we generated a custom RepeatModeler database, which led to the annotation of 30.09% of sequenced reads (S2 Table). The most abundant sequences derived from the genes *dph6* (22.51%) and *robo1* (3.43%) (Fig 3b). Among transposable elements (TEs), endogenous retroviruses (ERVs) constituted 3.97% of the total reads, representing one of the most prominent repeat groups in the GRC belt, particularly ERV1, ERVK, and ERVL. Other TEs, such as LINEs, SINEs, and DNA transposons, collectively accounted for less than 0.13%. We also detected trace amounts of RNAs (0.03%), satellite sequences (0.003%), and *COPS2*, *BMP15*, and *SECISBP2L* genes (each less than 0.0002%), likely arising from contamination by transcripts.

To confirm the identification of the major components of the GRC belts, we performed FISH with probes specific to the *dph6* gene and the most abundant ERVK, TguLTRK1. Both probes hybridized in the GRC belts (Fig 3c). Additionally, we localized the *dph6* probe on the metaphase GRC from testis it in the middle of the q-arm (S2a Fig), where the *dph6*-positive region corresponded to a bright DAPI-positive segment (S2b Fig). Hybridization of TguLTRK1 to metaphase chromosomes revealed that this element is an interspersed repeat widespread on many chromosomes, with a notable accumulation on the W chromosome and a pair of microchromosomes (S3 Fig).

### Mapping the Lampbrush GRC with Repetitive Sequence Probes

To test whether the GRC contains interstitial telomeric sequences, we performed FISH with a probe specific to the vertebrate telomeric repeat (TTAGGG)n. The telomeric probe produced signals exclusively at the chromosome ends of the GRC (S4 Fig), further supporting the hypothesis that the zebra finch GRC did not originate through recent fusion of microchromosomes [2]. Then, we tested whether the known zebra finch centromeric repeats *Tgut191A* and *Tgut716A* [20,41] could mark the GRC centromere. The hybridization with the *Tgut191A* probe did not yield any detectable signal on the GRC (S5a Fig), while mapping with a probe to *Tgut716A* satellite produced inconsistent results: dot signals were observed in one of the GRC belts, sometimes in both belts, or were completely absent (S5 Fig).

To further characterize the structural organization of the GRC, we identified new GRC-associated tandem repeats, *Tgut17-167* and *Tgut16-201* (S3 Table) and examined their distribution. The *Tgut16-201* satellite was detected in the assembly generated from PacBio HiFi testis sequencing [12]. Simulating an Illumina library from this assembly for RepeatExplorer analysis, we identified the *Tgut16-201* cluster and showed its enrichment in testis versus somatic Illumina reads. The satellite *Tgut17-167* was absent from the PacBio HiFi testis library, but was found in Illumina and Oxford Nanopore testis libraries [12]. We constructed a consensus sequence for *Tgut17-167* using a collection of ultra-long Nanopore reads and used the consensus sequences to design primers for PCR-generation of FISH probes. The probes to the *Tgut17-167* repeat, Sat167p1 and Sat167p2, were mapped on several chromomers near the chiasma distant from the belts (Fig 4a and 4b). The probe to *Tgut16-201*, Sat201, hybridised to the individual chromomeres on each GRC semi-bivalent just above the chiasma adjacent to the large coilin-containing body (Fig 4c).

**Figure 4.**
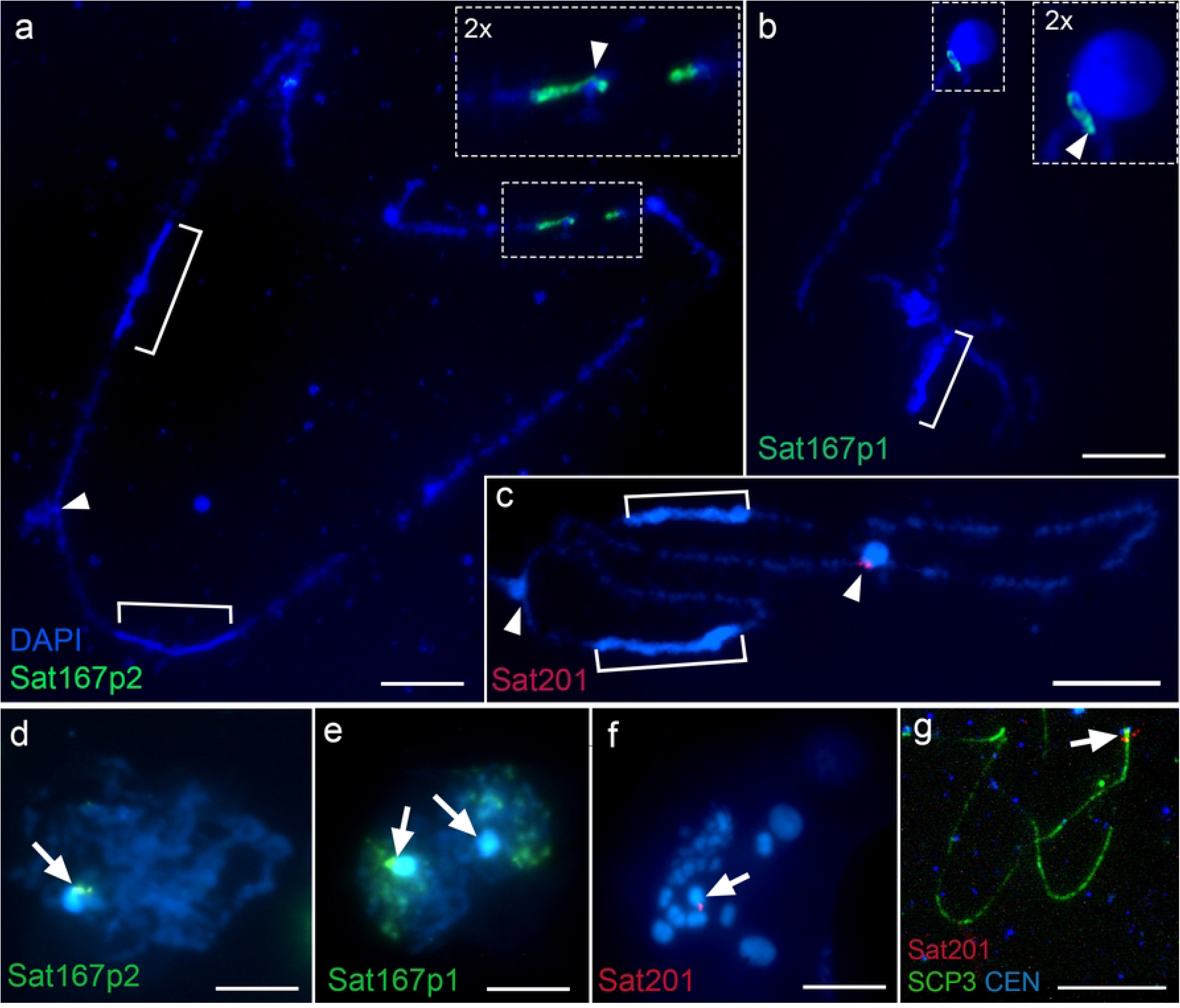
Localisation of Sat167p2, Sat167p1, and Sat201 probes, corresponding to *Tgut17-167* and *Tgut16-201* tandem repeats on the zebra finch GRC. FISH with (a) Sat167p2, (b) Sat167p1, and (c) Sat201 probes on the lampbrush GRC shows terminal signals near the chiasma (arrowheads) at the chromosome end opposite to the heterochromatic belts (square brackets). Localisation of Sat167p2 (d) and Sat167p1 (e) (green fluorescence) in spread spermatocyte nuclei. Arrows indicate a DAPI-bright heterochromatic body containing GRC material. (f) Localisation of the Sat201 probe on metaphase chromosomes from spermatocytes; the arrow indicates the GRC with a bright signal (red fluorescence). In images (a-f), chromosomes are counterstained with DAPI. (g) Localisation of the Sat201 probe (red fluorescence) on the synaptonemal complex of the pachytene GRC. The lateral elements of the synaptonemal complex are immunostained with antibodies against SCP3 (green fluorescence), while kinetochore and centromere proteins are detected using CREST antiserum (CEN, blue fluorescence). The arrow indicates the Sat201 probe signal adjacent to the functional centromere. Scale bars: 10 µm.

The specificity of the Sat167p1, Sat167p2 and Sat201 probes for the GRC was checked by FISH on testis cell preparations. As a result, FISH with probes for Sat201 and Sat167p2 produced bright single-point signals exclusively on the DAPI-positive GRC body (Fig 4d) or in the centromere region of the acrocentric metaphase GRC (Fig 4f and 6Sb Fig), whereas the Sat167p1 probe also hybridized to additional genomic loci (Fig 4e). To confirm the pericentromeric location of the *Tgut16-201* and *Tgut17-167* repeats, we hybridized the Sat201 probe on female synaptonemal complexes preparations combined with immunodetection of centromeric proteins. As a result, the GRC functional centromere was found to be terminally located, with the *Tgut16-201* locus adjacent to it (Fig 4g). A similar pattern was observed after hybridization with the Sat167p2 probe (S6a Fig).

### Cytogenetic Map of the Zebra Finch GRC

To integrate our findings with previously reported cytogenetic data on the zebra finch GRC, we constructed a cytological map of the lampbrush GRC based on its chromomere pattern and localised key repetitive elements on the map (Fig 5). Prior to this study, only two types of markers had been mapped on the GRC: lambda DNA clone 27L4 (GenBank accession number FJ609199.1) [42], and amplified fragments of the *dph6* gene [21]. We showed that GRC belts are highly enriched in specific ERVs (ERV1, ERVL, and ERVK) and *dph6*-derived sequences. The *dph6* region may comprise up to ∼50 Mb of the GRC, according to the latest alignment of the GRC to the main chromosome set in the zebra finch [12,43]. The 27L4 probe, prepared for FISH by nick-translation, hybridised along almost the entire length of the GRC (Fig 5b), leaving a gap in the middle [42]. The position of this gap seems to correspond to the region labeled by our microdissected GRC belt probe on the pachytene chromosome (Fig 5b and S1c Fig), which supports the observation that the belt regions and the remainder of the GRC may differ in their TE content.

**Figure 5.**
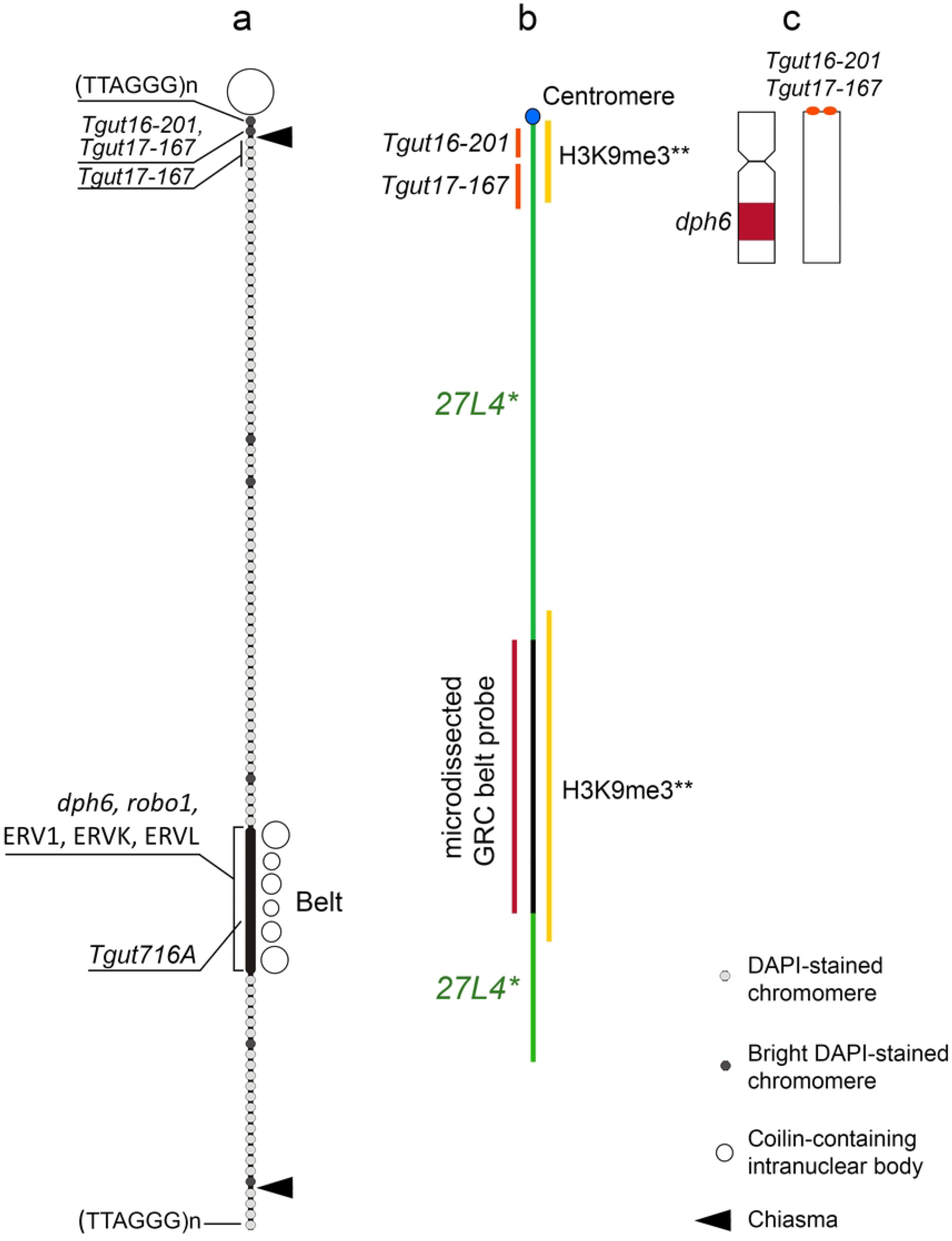
Cytogenetic maps of the zebra finch GRC (not to scale). (a) Lampbrush GRC map, showing the chromomere pattern, positions of coilin-containing NBioMCs, and the physical locations of (TTAGGG)n, *Tgut716*, *Tgut16-201*, *Tgut17-167*, *dph6,* and ERVK sequences (exemplified by the most abundant *TguLTRK1*), as well as sequences identified by bioinformatic analysis of the GRC belt region (*robo1*, ERV1 and ERVL). (b) Pachytene GRC map, with the location of the 27L4 probe from Itoh et al., 2009 [42] (27L4*) and the H3K9me3 enrichment pattern from Schoenmakers et al., 2010 [25] (H3K9me3**). (c) Metaphase GRC, observed in both submetacentric and acrocentric forms in preparations from dividing spermatocytes. The *dph6*-enriched belt corresponds to the DAPI-positive block in the middle of the q-arm of the submetacentric GRC, while the *Tgut16-201* and *Tgut17-167*,loci are located at the terminal end of the acrocentric GRC.

Another notable cytological marker of the female zebra finch GRC at the pachytene stage was shown by Schoenmakers et al. [25] – the enrichment of H3K9me3 at the terminal and middle parts in the GRC synaptonemal complex. These regions seem to correspond to the centromeric and belt regions of the GRC, respectively (Fig 5b). The location of MLH1 foci on pachytene chromosomes is also an important component of the cytogenetic map; two terminal MLH1 foci have been reported on the GRC synaptonemal complex spreads [2,14]. The localisation of chiasmata on the lampbrush GRC is in good agreement with these data.

## Discussion

### Centromeric features of GRC belts

The avian LBCs provide a powerful system for cytogenetic analysis, enabling accurate localisation of sequences and simultaneously assessing their transcriptional activity, yet, identifying such basic chromosome components as centromeres remains challenging. Immunostaining with antibodies against CENP-A, CENP-B, or CREST has not yielded reliable signals on avian LBCs [27], consequently, FISH using centromeric repeats has become the principal approach for mapping centromeres [36,44,45]. On LBCs, repeat-defined centromeric regions were shown to be associated with NBioMCs enriched in specific proteins, primarily components of the cohesion complex [40,44,46,47]. This reproducible association makes NBioMCs reliable cytological landmarks for centromere localisation on LBCs [27,48]. In the zebra finch, centromeric regions of autosomes and sex chromosomes are associated with coilin-containing NBioMCs [40]. Accordingly, the terminal functional centromere of the lampbrush GRC is associated with the prominent NBioMC, further supporting the use of NBioMCs as markers of centromeres in avian LBCs. Therefore, the presence of additional coilin-positive NBioMCs within the GRC belts together with the localisation of the canonical centromeric repeat *Tgut716A* in the same region suggests that the belts possess certain features of centromeric chromatin.

### A model for non-Mendelian GRC inheritance

The centromeric properties of the GRC belts may contribute to persistent chromatid cohesion and thereby explain the unusual behaviour of the chromosome during cell division. During early zebra finch embryogenesis, non-disjunction within the *dph6*-positive region, which corresponds to the GRC belts identified here, results in GRC lagging and subsequent elimination of the chromosome from somatic cells [49]. Thus, the GRC belts may have the ability to maintain prolonged sister chromatid cohesion, leading to the chromosome non-disjunction during anaphase. We hypothesise that a similar mechanism may also operate during female meiosis II, promoting sister chromatid non-disjunction and ensuring that both sister chromatids of the same GRC are transmitted into the egg nucleus. The molecular basis of this persistent cohesion remains unknown. Possible mechanisms include failure to assemble a functional kinetochore within the GRC belts, defective removal of cohesin complexes, or the persistence of DNA catenanes that cannot be resolved by topoisomerase II. Distinguishing among these possibilities will require further investigation of GRC composition and behaviour after the lampbrush stage and throughout meiotic divisions, which remains technically challenging in avian systems.

Although speculative, this model resembles the behaviour of the X chromosome in the fungus gnat *Bradysia coprophila* [50]. In this species, *cis*-acting controlling elements regulate sister chromatid non-disjunction during male meiosis II by preventing kinetochore formation and proper alignment of the X chromosome on the metaphase plate. As a result, both sister chromatids of the X chromosome segregate into a single spermatid, which subsequently develops into the functional sperm [50–52]. During embryogenesis, chromatid non-disjunction likewise leads to the elimination of X chromosomes. Although the molecular nature of the *B. coprophila* controlling element remains unknown, its translocation to autosomes has been shown to induce non-disjunction and subsequent chromosome elimination in a *cis*-acting manner [50].

If a similar controlling element acts in the zebra finch GRC inheritance, the repeated transmission of two identical GRC sister chromatids through the female germline could ultimately contribute to the exceptionally low nucleotide diversity, reported for this chromosome by Chen et al. [43]. A similar effect can be obtained if the GRC segregates normally during female meiosis, resulting in the egg receiving a single GRC chromatid, and if the nondisjunction of the GRC occurs later during embryogenesis in female primordial germ cells, causing bivalent formation in the oocyte. Thus, whether the GRC belts affect two-chromatid transmission in female meiosis or cause chromosome nondisjunction in primordial germ cells remains to be revealed in future studies.

### Unique architecture of passerine acrocentric GRCs

Using high-resolution mapping on the lampbrush GRC, we showed the functional terminally located centromere lacks the canonical zebra finch centromeric repeats *Tgut191A* and *Tgut716A* and instead contains the GRC-specific tandem repeats *Tgut17-167* and *Tgut16-201.* This unusual centromeric composition is consistent with observations in other passerines, two nightingale species *Luscinia megarhynchos* and *L. luscinia*, in which the GRC centromere similarly lacks the major genomic centromeric repeats and is enriched in GRC-specific satellite sequences [53].

Notably, the acrocentric morphology of the GRC has been documented not only in the zebra finch females [1,14,25,42], but also in all other passerine species studied to date [2,5,16,17]. In these species, the GRC bivalent consistently carries a chiasma adjacent to the centromere, despite recombination typically being suppressed in centromeric regions [54,55]. Such a persistent pericentromeric chiasma may facilitate the rapid accumulation of GRC-specific satellite repeats, observed in the zebra finch and nightingales, through unequal crossing over. More interestingly, proximal chiasmata have been reported to promote chromosome missegregation during meiosis I and II, in contrast to distal crossing overs, which generally ensure accurate segregation of homologous chromosomes [56–59]. However, the distinctive properties of the avian oocyte spindle, including its asymmetric architecture, delayed chromosome congression, and tolerance of atypical kinetochore–microtubule interactions [60], may mitigate the destabilising effects of proximal chiasmata.

## Conclusion

The study of the lampbrush GRC highlights the interplay between centromere position, crossing over (chiasma) site, and centromeric sequence composition in determining chromosome fate. Our data reveal an unusual architecture of the zebra finch GRC comprising two distinct centromere-associated domains: an evolutionary young functional terminal centromere enriched in GRC-specific repeats and a heterochromatic belt region containing canonical centromere sequence. We propose that interaction between these domains, together with pericentromeric chiasma, may contribute to the unique inheritance dynamics of the GRC, promoting its transmission through the female germline while facilitating its elimination from somatic lineages during early embryogenesis.

## Material and Methods

All procedures involving animals were conducted in accordance with the protocols approved by the Ethical Committee of Saint Petersburg State University (#131-03-3, issued on 01.06.2017, #131-03-4, issued on 13.03.2024) and were performed following relevant guidelines and regulations. We use two male and ten female zebra finches, as well as a five-day-old chick and two embryos. All birds were obtained from the breeding colony maintained at the Saint Petersburg State University facility.

### Lampbrush Chromosome Preparations

LBCs were manually isolated from previtellogenic or early vitellogenic oocytes (0.3-1 mm in diameter) following a standard microsurgical technique for germinal vesicle isolation [61]. Preparations were fixed in 2% paraformaldehyde (Sigma-Aldrich, USA), dehydrated in absolute ethanol, air-dried, and stored frozen. For immunostaining, preparations were kept in 70% ethanol at +4°C and used within one week. For microdissection, they were fixed exclusively in a series of 50% and 70% ethanol, omitting both paraformaldehyde fixation and dehydration with 96% ethanol. These preparations were stored in 70% ethanol until use and air-dried immediately before use.

### Meiotic Chromosome Preparation from Testes

Meiotic chromosomes from zebra finch testes were prepared according to the method by MacGregor and Varley [62]. Briefly, testes from adult birds were dissected in a 2.3% sodium citrate solution (Helikon, Russia). The testis capsule was removed, and the parenchymal tissue was minced. The cell suspension was transferred to a new tube and incubated for 15 minutes at room temperature. Cells were then pelleted by centrifugation at 800 rpm for 10 minutes (Eppendorf, Germany) and resuspended in an ice-cold fixative solution of methanol and glacial acetic acid (3:1). After 15 minutes of fixation, the cells were pelleted again and resuspended in fresh fixative. The fixed cell suspension was stored at -20°C for 2 hours to overnight. Chromosome preparations were made by dropping the fixed cell suspension onto pre-cooled, wet slides.

### Preparation of Synaptonemal Complexes

Chromosome spreads from one-week-old chick ovaries were prepared using the drying-down method [63]. Ovary was incubated in an extraction buffer (30 mM Tris, 50 mM sucrose, 17 mM trisodium citrate dihydrate, 5 mM EDTA, pH 8.2; all components from Helicon, Russia) for 30-60 minutes. Ovary was then macerated on a glass slide in 100 µl of 100 mM sucrose (pH 8.2). Subsequently, 20 µl aliquots of the resulting cell suspension were gently dropped onto the corner of a new slide pre-coated with a solution of 1% paraformaldehyde and 0.15% Triton X-100 (Applichem, Germany), pH 9.2. Using a seesaw motion, the suspension was spread across the slide surface. The slides were then incubated in a humid chamber for 1 hour, rinsed in 0.4% Photo-Flo (Kodak, USA), and air-dried. Preparations were stored at -20°C until use.

### Immunofluorescent Staining

Lampbrush chromosome preparations were rehydrated through a series of 5-minute incubations in 50% ethanol, 30% ethanol, and phosphate-buffered saline (PBS; 137 mM NaCl, 2.7 mM KCl, 10 mM Na₂HPO₄, 1.8 mM KH₂PO₄, pH 7.4; all components from Helicon, Russia). To minimise nonspecific antibody binding, slides were blocked with 1% bovine serum albumin (BSA; Thermo Fisher Scientific, USA) in PBS for 30 minutes at 37°C. Both primary and secondary antibodies were diluted in 1% BSA/PBS solution according to manufacturer’s recommendations. For coilin detection, slides were incubated overnight at 4°C with a primary rabbit anti-coilin antibody (Thermo Fisher Scientific, USA Cat# PA5-110950, RRID:AB_2856361, 1:400), followed by a 40-minute incubation at 37°C with a secondary goat anti-rabbit antibody conjugated to Alexa Fluor 594 (Thermo Fisher Scientific, USA). For 5-methylcytosine (5mC) detection, an antigen demasking step was required prior to blocking: slides were incubated in 2M HCl for 1 hour and washed three times for 5 minutes each in PBS. After this pretreatment, the staining procedure was identical to that used for coilin, using a primary rabbit anti-5mC antibody (Abcam, UK Cat# RM231, RRID: ab214727, 1:400) incubated for 1 hour at 37°C. Following each antibody incubation, slides were washed three times in PBS containing 0.1% Tween 20 (MP Biomedicals, USA).

Centromeres and synaptonemal complexes on chromosome spreads were visualised using human anti-centromere (Antibodies Inc., USA) and rabbit anti-SCP3 (Thermo Fisher Scientific, USA) antibodies, following a protocol adapted from Malinovskaya et al. [8]. Slides were permeabilised in 0.1% Triton X-100 (AppliChem GmbH, Germany) in PBS for 45 minutes, subjected to antigen retrieval in 10 mM sodium citrate buffer (pH ∼6.0) at 80°C for 8 minutes, and then blocked with 1% BSA in PBS for 40 minutes at room temperature. Subsequently, slides were incubated with primary antibodies overnight at 37°C. Following primary incubation, preparations were washed three times for 5 minutes each in PBS and incubated with secondary antibodies for 40 minutes at 37°C: anti-human IgG conjugated to Cy3 (PA1-31226, 1:200) and anti-rabbit IgG conjugated to Alexa Fluor 488 (Thermo Fisher Scientific, USA, Cat# A27033, RRID: AB_2536096, 1:200). A final wash series was performed as described above.

All preparations were counterstained with 1 µg/mL DAPI (4′,6-diamidino-2-phenylindole; Polysciences, USA) in an antifade mounting medium based on DABCO (1,4-diazabicyclo[2.2.2]octane; Sigma-Aldrich, USA) in PBS and 50% glycerol (Sigma, USA).

### Microdissection and Whole-Genome Amplification

GRC loopless belts were microdissected using an AXIOVERT 10 inverted microscope (Zeiss, Germany) equipped with a micromanipulator (Zeiss MR mot) and a mechanical positioner. Microdissection needles were prepared from 2 mm-diameter glass rods (Duranglass, Germany) using a pipette puller (Narishige, PB-7, Japan). Glass Pasteur micropipettes, used as collection tubes, were pre-treated by silanisation with a 1% solution of dimethyldichlorosilane in carbon tetrachloride, rinsed three times with 1 mM EDTA (pH 7.5; Helicon, Russia), dried at 60°C for 30 minutes, baked at 100°C for 1 hour, and sterilised by UV irradiation overnight. Following microdissection, the needle tip containing the isolated chromosome fragment was broken off into a pre-treated collection tube. The microdissected material was deproteinised in a solution containing 0.5 mg/mL proteinase K (Sigma, USA), 10 mM Tris-HCl (pH 7.5), 10 mM NaCl, 0.1% Triton X-100, and 30% glycerol (all components from Helicon, Russia) at room temperature. In total, chromosomal material from 7.5 GRC bivalents (equivalent to 30 chromatids) was collected. For further sequencing and probe preparation, the microdissected DNA was amplified using the GenomePlex Single Cell Whole Genome Amplification Kit (WGA4, Sigma-Aldrich, USA) according to the manufacturer’s instructions. Amplification efficiency was assessed by DNA electrophoresis in a 1% agarose gel. Reactions were considered successful if they yielded a visible smear of DNA fragments ranging from 200 to 1000 bp at a minimum concentration of 100 ng/μL.

### Sequencing and Bioinformatic Analysis

The DNA library was prepared using the NEBNext Ultra DNA Library Prep Kit (New England Biolabs, USA) and sequenced on an Illumina NextSeq 550 platform in single-end read mode. The quality of the raw reads was assessed using FastQC (www.bioinformatics.babraham.ac.uk/projects/fastqc). Subsequent processing and analysis were conducted in Geneious 9.0 software (https://www.geneious.com), which involved the removal of protozoan contaminants and poly-G artifacts. Then reads were trimmed [64] and the obtained library (uploaded in NCBI: SRR38807074) was aligned to the zebra finch reference genome (bTaeGut1.4.pri, NCBI accession number GCF_003957565.2). *De novo* repeat identification was performed using a repeat database for the GRC assembly generated with RepeatModeler [65]. We identified repeats from the highly amplified regions in the GRC, i.e., the *dph6* regions as well as the one from chromosome 1 including *robo1* and *gbe1* and the surrounding genes. For this we performed a RepeatMasker search with the *dph6* and the *robo1* genes. The analysis utilised a custom repeat database constructed from the following sources: the GRC HiFi assembly [43], avian repeats from Repbase (Genetic Information Research Institute) [66], and known zebra finch satellite sequences [20].

### GRC Satellite Identification

To characterize new FISH markers for the GRC zebra finch, we used two strategies. First, we used the zebra finch GRC assembly generated from PacBio HiFi testis reads from Chen et al. [43]. Then, we simulated an Illumina library from the GRC assembly with the “Get pseudo short paired end reads from long reads” tool from the RepeatExplorer’s Galaxy (https://repeatexplorer-elixir.cerit-sc.cz/galaxy/) and prepared the reads to be used as an input for RepeatExplorer. Afterwards, we performed RepeatMasker alignments of the soma and testis Illumina reads from Kinsella et al. [21] against the RepeatExplorer contigs to identify candidates satellite clusters with a clear enrichment in the testis sample versus the somatic one. This way, we detected the satellite *Tgut16-201 ()*. We generated primers in the primer3 software [67] using the consensus sequence of this selected satellite.

We also applied a complementary approach to identify germline-enriched satellites. For this, we run TAREAN [68] to publicly available testis and somatic Illumina libraries, with both tissues sampled from the same zebra finch individuals [15]. The comparative analysis between soma and testis libraries allowed the discovery of a testis-enriched satellite *Tgut17-167* in the testis library SRX11324199. To validate the testis-specificity of the satellite, we also ran Repeatmasker (V4.1.0) using the TAREAN consensus sequence for *Tgut17-167* on both liver (SRX11324214) and testis (SRX11324199) tissues of the zebra finch male (BR19117). Next, we used the divergence summary table generated by Repeatmasker (divsum output file) to plot a subtractive landscape of the satellite distribution in the liver and testis libraries using ggplot2 (V3.5.1) [69]. To further investigate the organization of the satellite, we also scanned long and ultra-long reads for the identification of satellite arrays. For this, we used two testis libraries generated by PacBio HiFi sequencing and Oxford Nanopore sequencing [12], respectively, and ran Repeatmasker using the consensus sequence produced by TAREAN (*Tgut17-167*). Hits were solely detected in the Nanopore library and were subsequently extracted. A collection of ultra-long Nanopore reads spanned by the satellite array was obtained and used as a query sequence to run Tandem Repeat Finder (TRF) [70]. Since monomers of this satellite were very divergent, we designed primers by manually cutting monomers of a Nanopore read and aligning them within the Geneious software. Then, we identified conserved regions and designed two primer pairs anchoring in these regions with primer3 [67].

### Probes Generation and FISH

A biotin-labelled oligonucleotide probe C(TAACCC)_5_TAAC corresponding to the telomeric repeat was generated using an ASM-800 DNA/RNA synthesiser (Biosset, Russia). Probes for the *Tgut16-201* and *Tgut17-167* repeats, centromere-specific repeats *Tgut716A* and *Tgut191A,* the interspersed repeat *TguLTRK1* (which refers to the TE0 repeat [41]), and *dph6 (*dph6-611 and dph6-801) [21] were generated by PCR with specific primers (S3 Table). Zebra finch genomic DNA from somatic tissues was used as the PCR template for Tgut716A, Tgut191A, and TguLTRK1 probes, testis tissue for dph6-611 and dph6-801 probes, and microdissected GRC loopless region material for the GRC belt-specific probe. The PCR mixture contained 1x Taq buffer, 2.5 mM MgCl_2_, 0.2 mM of each dNTP, 10 μM of each forward and reverse primer, 2.5 units of Taq DNA polymerase (all components from Sileks, Russia), and 20 ng of template DNA. For the generation of labelled probes, 0.04 mM dTTP in the reaction mixture was replaced with an equimolar concentration of biotin-16-dUTP (Sileks, Russia) or digoxigenin-11-dUTP (Sigma-Aldrich, USA).

Labelled probes were diluted to a final concentration of 20 ng/μL each in a hybridisation buffer containing 50% formamide, 10% dextran sulphate, 2xSSC, and a 50-fold excess of *E. coli* tRNA (all components from Sigma-Aldrich, USA). FISH on LBCs was performed according to the DNA/DNA+RNA hybridisation protocol without pretreatment, as described previously [29,34]. Hybridisation was carried out overnight at room temperature for the telomeric probe and at 37°C for all other probes. Biotin-labelled probes were detected using avidin-FITC and anti-avidin-FITC (Sigma-Aldrich, USA). To enhance the signal for centromeric and telomeric probes a Tyramide Amplification Kit with Alexa488 (ThermoFisher, USA) was used. Digoxigenin-labelled probes were detected using anti-digoxigenin-Cy3 antibodies (ThermoFisher Scientific, USA). Finally, the slides were dehydrated in 70% and 96% ethanol, air-dried, and mounted in an antifade solution consisting of 50% glycerol, 2xSSC, 1% DABCO, and 5 μg/mL DAPI.

The FISH procedure on meiotic chromosomes from testis and on chromosome spreads following immunocytochemistry was similar to the standard protocol with some modifications. Prior to hybridisation, preparations were treated with 1 U of RiboShroeder (JenaBioscience, Germany) in 2xSSC for 1 hour at 37°C, followed by three washes in 2xSSC for 5 minutes. Additionally, meiotic chromosomes from testis were treated with pepsin as described by Liehr et al. [71].

### Microscopy and Cytological Map Construction

Chromosome preparations were examined using a Leica DM4000B fluorescence microscope equipped with a black-and-white CCD camera and appropriate filter cubes (Leica Microsystems, Germany). Image acquisition and processing were performed using LAS AF software (Leica Microsystems, Germany). For the construction of the cytological map, we analysed 20 images of well-spread GRC bivalents isolated from the oocytes of comparable size of seven sexually mature female zebra finches. The absolute length of each GRC and the relative positions of cytological markers were measured using ImageJ software [72]. The final cytological map was drawn using CorelDRAW Graphics Suite 2017 and figure assembly was performed using Adobe Photoshop CS5 (Adobe Systems).

## Acknowledgements

We thank Lyubov Malinovskaya, Pavel Borodin, and Anna Pendina for kindly providing antibodies. We are grateful to the “Chromas” Resource Centre and the Centre for Molecular and Cell Technologies at the Saint Petersburg State University Scientific Park for access to their equipment. Sequencing was carried out at the Genomics Resource Centre of the Institute of Cytology and Genetics, SB RAS (Novosibirsk).

## Statement of Ethics

All applicable international, national, institutional guidelines for the care and use of animals were followed. All experiments were approved by the Ethics Committee on Animal Research of Saint Petersburg State University, Russia (#131-03-3, issued on 01.06.2017, #131-03-4, issued on 13.03.2024).

## Authors’ Contributions

O.T., S.G. and V.V. made experiments and interpreted the results. J.J. constructed cytological maps. N.R. and K.Z. made microdissection and generated probes for the GRC belts. N.V., F.R., and A.S. made bioinformatic analysis. M.K. and S.G. made chromosome mapping. O.T., E.G., and S.G. wrote a manuscript draft. S.G. conceived and designed the study. All authors read and approved the final manuscript.

## Data Availability Statement

DNA sequencing data have been submitted to the NCBI SRA (accession number: SRR38807074). Other data are within the text and its supporting information.

